# MATEdb2, a collection of high-quality metazoan proteomes across the Animal Tree of Life to speed up phylogenomic studies

**DOI:** 10.1101/2024.02.21.581367

**Authors:** Gemma I. Martínez-Redondo, Carlos Vargas-Chávez, Klara Eleftheriadi, Lisandra Benítez-Álvarez, Marçal Vázquez-Valls, Rosa Fernández

## Abstract

Recent advances in high throughput sequencing have exponentially increased the number of genomic data available for animals (Metazoa) in the last decades, with high-quality chromosome-level genomes being published almost daily. Nevertheless, generating a new genome is not an easy task due to the high cost of genome sequencing, the high complexity of assembly, and the lack of standardized protocols for genome annotation. The lack of consensus in the annotation and publication of genome files hinders research by making researchers lose time in reformatting the files for their purposes but can also reduce the quality of the genetic repertoire for an evolutionary study. Thus, the use of transcriptomes obtained using the same pipeline as a proxy for the genetic content of species remains a valuable resource that is easier to obtain, cheaper, and more comparable than genomes. In a previous study, we presented the Metazoan Assemblies from Transcriptomic Ensembles database (MATEdb), a repository of high-quality transcriptomic and genomic data for the two most diverse animal phyla, Arthropoda and Mollusca. Here, we present the newest version of MATEdb (MATEdb2) that overcomes some of the previous limitations of our database: (1) we include data from all animal phyla where public data is available, (2) we provide gene annotations extracted from the original GFF genome files using the same pipeline. In total, we provide proteomes inferred from high-quality transcriptomic or genomic data for almost 1000 animal species, including the longest isoforms, all isoforms, and functional annotation based on sequence homology and protein language models, as well as the embedding representations of the sequences. We believe this new version of MATEdb will accelerate research on animal phylogenomics while saving thousands of hours of computational work in a plea for open, greener, and collaborative science.

## Introduction

In the midst of an explosion in the availability of genomic sequences, the advancement of phylogenomic, phylotranscriptomic, and comparative genomic studies in animals is hindered by the preprocessing and homogenization of the input data. With high-quality chromosome-level genomes being published almost daily in the last few years, we are gaining access to new biological knowledge that is helping to solve trickier scientific questions, such as the identity of the sister taxon to Bivalvia (Song et al., 2023) or the evolution of non-coding and repetitive regions (Osmanski et al., 2023). In addition, the use of transcriptomes as a proxy of a species proteome continues to be a main source of proteome data as a cheaper and easier alternative for phylogenetic inference (Erséus et al., 2020; Mongiardino Koch et al., 2018; Zapata et al., 2014, among others) and gene repertoire evolution (De Oliveira et al., 2016; Fernández & Gabaldón, 2020; Thoma et al., 2019) in less-studied animals.

Together, these genomic and transcriptomic studies have provided a vast number of resources for a plethora of animals that cannot be directly used in phylogenomic studies before proper preprocessing. This is especially true for older datasets where data quality is much lower and can have a high impact on the results obtained. Moreover, the use of different computational pipelines for data processing makes data not comparable and prone to false positives and negatives. For instance, the transcriptome assembly methodology used can impact the comparability among different datasets (e.g. for the subset of mollusk transcriptomes obtained from Krug et al. (2022), the number of ‘genes’ inferred with Trinity is significantly different -p-value < 0.1 - to the ones we obtained, Figure S1), while the ‘ready-to-use’ protein files provided in some genome sequencing projects cannot be easily matched with the other genome files for additional analyses due to different nomenclature across files. This mainly impacts research groups with lower computational resources or experience who cannot leverage the publicly available data into their research. To help alleviate these issues, we previously published the Metazoan Assemblies from Transcriptomic Ensembles database (MATEdb) containing high-quality transcriptome assemblies for 335 arthropods and mollusks (Fernández et al., 2022). Here, we present its second version, MATEdb2, that differs from the previous one in three main aspects: (1) we have increased the taxonomic sampling to all animal phyla with high-quality data publicly available, and provide the first transcriptomic sequences for some animal taxa; (2) we include a standardized pipeline for obtaining the protein sequences from GFF genomic files instead of adding the precomputed protein files publicly available, making it easier to replicate and combine with the associated genomic sequence; (3) we provide the functional annotation of all proteins using a language-based new methodology that outperforms traditional methods (Barrios-Núñez et al., 2024). We hope that this newer version of MATEdb accelerates research on animal evolution by providing a wider taxonomic resource of high-quality proteomes across the Animal Tree of Life.

## Material and Methods

### Increased taxonomic coverage

The first version of MATEdb (Fernández et al., 2022) included high-quality datasets from 335 species of arthropods and mollusks, with special attention to lineage representation within each phylum. Here, we provide a newer version of MATEdb that expands the taxonomic representation across the Animal Tree of Life by incorporating a total of 970 species from virtually all animal phyla that have publicly available genomic or transcriptomic data, as well as some outgroup species relevant for understanding animal evolution. Taxon sampling tried to maximize the taxonomic representation within each phylum while considering the quality of the data. The number of proteomes per phylum included in MATEdb2 and the distribution of gene number are represented in Figures 1a and b, and the complete list of species and their metadata is included in Table S1.

**Figure 1.**
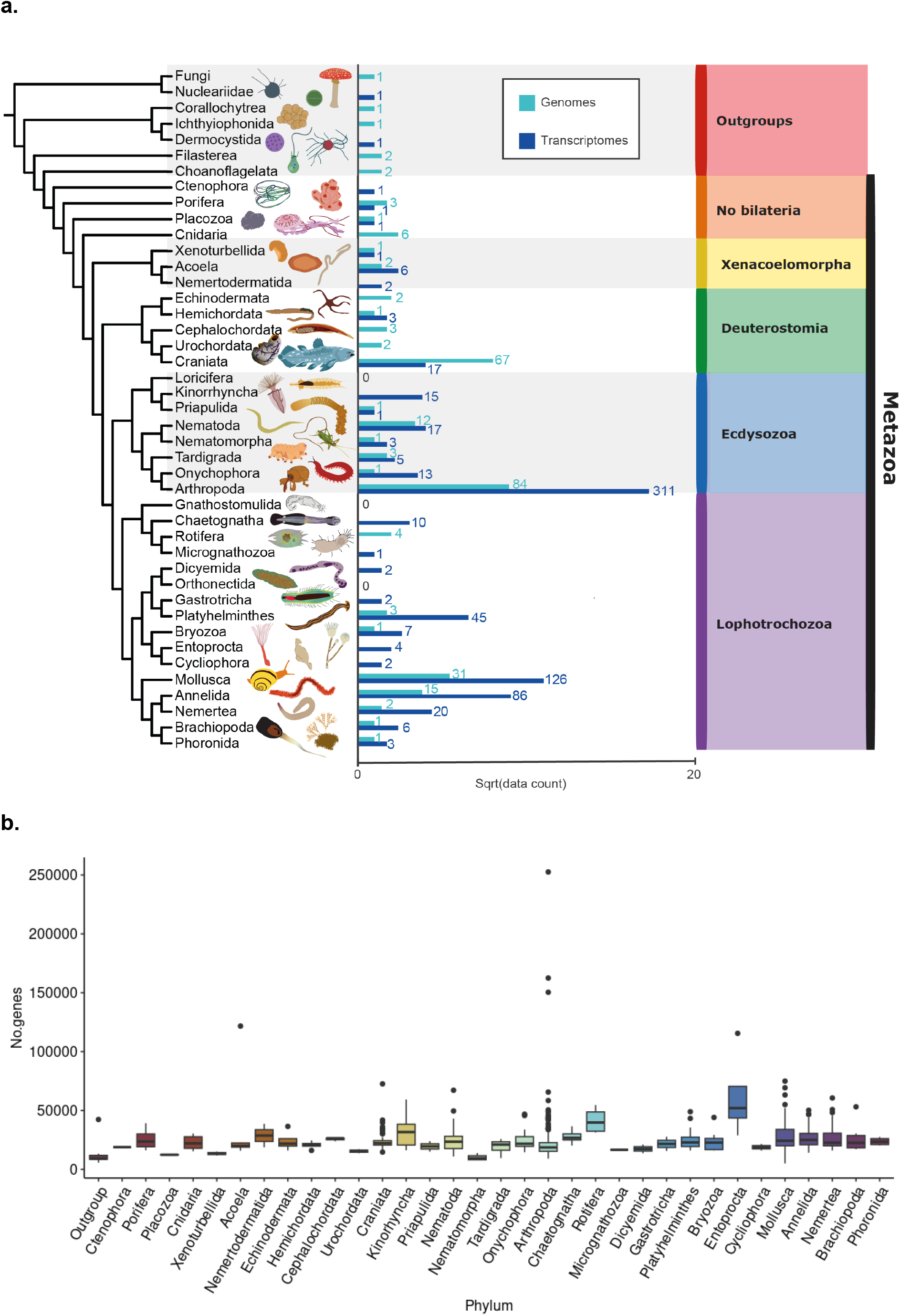
Taxonomic representation of species included in MATEdb2. **a**. The number of datasets per phylum separated by the type of data (genomes and transcriptomes in light and dark blue, respectively). **b**. Distribution of the number of genes per proteome across phyla.

### Improved analytical pipeline for genomes

In the previous version of MATEdb (Fernández et al., 2022), we directly downloaded the Coding DNA Sequences (CDS) and proteome files from the public repositories in the case of genomes. However, a closer inspection of both files together with their corresponding genome sequence and annotation revealed incongruences between them that needed to be manually curated. Just looking at gene numbers, only 9 out of the 59 genomes we are keeping from the previous version of MATEdb had the same number of protein-coding genes in the peptide and GFF files. In addition, most of the proteomes differed in more than 1000 genes, with 15 having more than 10k of difference. This is caused by the lack of consensus in the annotation and publication of genome files, with some authors uploading modified versions of the protein sequences that do not map directly with the reported GFF and FASTA file, hindering the utility of those files for additional analyses. Another case we encountered when building the new version MATEdb2 was for the only chromosome-level ctenophore genome available back then (*Hormiphora californensis*), in which the proteins extracted directly from the GFF and FASTA file contained premature stop codons in virtually all the sequences, which made us discard this species. Moreover, even highly curated public databases can contain wrong or missing data, such as the case of *Apis mellifera* and *Anopheles gambiae*’s CDS file in Uniprot (UniProt Consortium, 2023) containing only a couple of sequences instead of the whole proteome. Therefore, we have included in the newer version of MATEdb a standardized pipeline for obtaining the CDS and protein files using directly the FASTA and GFF files of the corresponding genome.

The analytical pipeline of MATEdb2 is shown in Figure 2. In brief, the differences with the pipeline depicted in MATEdb (Fernández et al., 2022) are the following: (1) we included a standardized pipeline for obtaining the longest isoform from genomes; (2) for a few exceptions, we lowered the threshold used to consider a dataset as high-quality to 70% C+F (complete plus fragmented) BUSCO score (Manni et al., 2021), as the original 85% threshold was too restrictive when prioritizing a wide taxonomic sampling and the inclusion of biologically interesting species that are not widely studied. Further details about the pipeline are shown in Figure 2.

**Figure 2.**
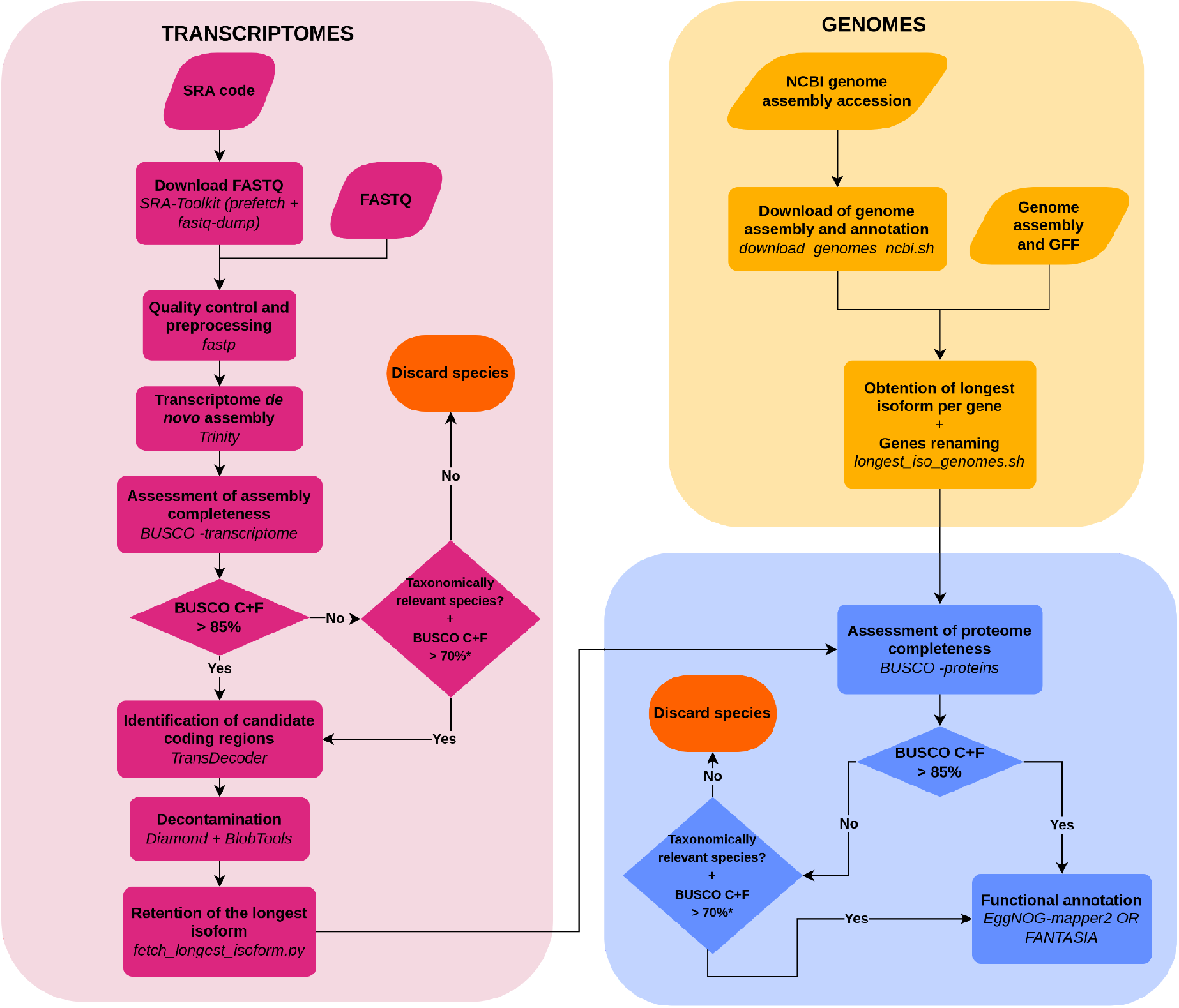
Pipeline followed to generate the MATEdb2 database. All steps differing from the MATEdb original pipeline are discussed in detail in the main text.

### Compilation of genomic data

Genome assembly (FASTA) and annotation (GFF) files for each species were downloaded through NCBI Datasets (Sayers et al., 2023) or from the direct URL download link for genomes available in other repositories. The database source for each species is referenced in the Supplementary Material, Table S1, while the bash script “download_genomes.sh” used to automatize the downloading of several files is included in the GitHub repository and the Singularity container (See Data Availability).

Once downloaded, we used AGAT (Dainat et al., 2023) to obtain the GFF containing only the longest isoforms which was then used to get the FASTA file with the longest protein sequence for each gene (and its corresponding CDS). In addition, we renamed the sequences to match the structure used in the transcriptomic part of the MATEdb2 pipeline and obtained a conversion file to keep track of the original names. These steps were performed using a custom bash script “longest_iso_genomes.sh”, also included in the GitHub repository and container.

Finally, gene completeness was assessed using BUSCO in protein mode against the metazoa_odb10 reference set (except for the outgroup species, where eukaryota_odb10 was used). More than 75% of our species passed the threshold of 85% complete plus fragmented used in MATEdb (Fernández et al., 2022). The remaining 25% includes almost all representatives of tardigrades, annelids, nematodes, acoels, and some representatives of other phyla (see Table S1). As we want to maximize the taxon representation of animal lineages while keeping datasets of high quality, we lowered the threshold value to 70% in these cases, a value previously used in other studies. These values may represent biological features of the genomes of these lineages (Barreira et al., 2021) or just a lack of representation of the lineage in the BUSCO reference datasets. As an exception, after this new threshold, 8 animal and 2 outgroup transcriptome assemblies have been included with a slightly lower BUSCO score due to their taxonomic relevance (e.g., they were one of the only two representatives of their lineage, such as in the case of the Priapulida). A list of discarded datasets can also be found in Table S2.

### Functional annotation of the gene repertoire

The longest isoform gene list for each dataset was annotated with the homology-based software eggNOG-mapper v2 (Cantalapiedra et al., 2021) and the FANTASIA pipeline (https://github.com/MetazoaPhylogenomicsLab/FANTASIA). FANTASIA is a pipeline that allows the annotation of whole proteomes using GOPredSim (Littmann et al., 2021), a protein language-based method that transfers GO terms based on embedding similarity. In brief, embeddings are vectorized representations of protein sequences generated using protein language models, such as ProtT5 (Elnaggar et al., 2022), that consider protein sequences as sentences and apply natural language processing tools to extract information from them. Here, besides the GO terms predicted by FANTASIA, we provide the raw per-protein ProtT5 embeddings. More details about the pipeline, the method or the benchmarking and comparison with homology-based methods can be checked elsewhere (Barrios-Núñez et al., 2024; Martínez-Redondo et al. 2024).

## Database availability

### Scripts and commands

Scripts and commands in the pipeline and the supplementary data (Tables S1-2, Figure S1) can be found in the following repository: https://github.com/MetazoaPhylogenomicsLab/MATEdb2

### Files deposited in the repository

For transcriptomes, the data repository contains (1) de novo transcriptome assemblies, (2) their candidate coding regions within transcripts (both at the level of nucleotide and amino acid sequences), (3) the coding regions filtered using their contamination profile (ie, only metazoan content or eukaryote for outgroups), (4) the longest isoforms of the amino acid candidate coding regions, (5) the gene content completeness score as assessed against the BUSCO reference sets, and (6) orthology and protein language-based gene annotations, and per-protein ProtT5 embeddings. In the case of genomes, only files (4), (5), and (6) are provided in MATEdb2, together with a filtered version of the file (3) with just the longest CDS per gene.

The database is hosted on our own server and will be there indefinitely. The database will be expanded as we incorporate new datasets from underrepresented lineages, such as nematodes, or as requested to be incorporated by the scientific community if resources allow it. Links for downloading can be found in the following file in the GitHub repository: https://github.com/MetazoaPhylogenomicsLab/MATEdb2/blob/main/linksforMATEdb2.txt

### Software availability

We provide a Singularity container for easy implementation of the tools used to generate the files in the database with the appropriate software versions along with their dependencies (https://cloud.sylabs.io/library/klarael.metazomics/matedb2/matedb2.sif). The software included is the following: SRA Toolkit v2.10.7 (http://ncbi.github.io/sra-tools/), fastp v0.20.1 (https://github.com/OpenGene/fastp; Chen et al., 2018), Trinity v2.11.0 (Grabherr et al. 2011), BUSCO v5.3.2 (Manni et al., 2021), TransDecoder v5.5.0 (https://github.com/TransDecoder/TransDecoder), Diamond v2.0.8 (Buchfink, Xie, & Huson, 2015), BlobTools v2.3.3 (Challis et al., 2020)., NCBI datasets v13.42.0, eggNOG-mapper v2.1.9 (Cantalapiedra et al., 2021), seqkit v2.1.0 (Shen, Sipos, and Zhao, 2024), AGAT v0.9.1 (Daniat et al., 2023), as well as some custom scripts.

## Discussion

We presented here the second version of MATEdb (MATEdb2), with almost 1000 animal species data. This newer version overcomes some of the previous restrictions of our database, including the restricted taxonomic representation of only arthropods and mollusks, and the use of previously preprocessed peptide and CDS files for genomes. Nevertheless, it is not devoid of limitations. The main limitation of this newer version are still the genome annotations. Even though we use an alternative approach that considers the incongruences found in some of the publicly preprocessed files, there may still be biases between the proteomes. These biases are caused by the heterogeneity of genome annotation methodologies, which can affect downstream analyses, such as ortholog inference (Weisman et al., 2022). These biases are typically ignored by phylogenomic studies that use publicly available preprocessed files. Nevertheless, correcting for this limitation by re-annotating the genomes using the same methodology is computationally expensive and is still biased toward species where additional data that improves this annotation (e.g. RNA-seq) is available.

## Author contributions

This database results from the collaborative effort of lab members from the Metazoa Phylogenomics Lab to offer the scientific community the possibility to reuse some of the data generated for their projects. GIMR, CVC, KE, LBA, and MVV contributed assemblies to the data repository. GIMR created the pipeline custom scripts for the genome data analyses and designed the MATEdb logo. KE created the Singularity container. CVC and RF contributed to the creation and management of the database. CVC created and curated the Github repository. RF provided resources and supervised the project. GIMR wrote the first version of the manuscript. All authors revised and approved the final version of the manuscript.

## Acknowledgments

GIMR acknowledges the support of Secretaria d’Universitats i Recerca del Departament d’Empresa i Coneixement de la Generalitat de Catalunya and ESF Investing in your future (grant 2021 FI_B 00476). RF acknowledges support from the following sources of funding: Ramón y Cajal fellowship (grant agreement no. RYC2017-22492 funded by MCIN/AEI /10.13039/501100011033 and ESF ‘Investing in your future’), the Agencia Estatal de Investigación (project PID2019-108824GA-I00 funded by MCIN/AEI/10.13039/501100011033), the European Research Council (this project has received funding from the European Research Council (ERC) under the European’s Union’s Horizon 2020 research and innovation programme (grant agreement no. 948281)), the Human Frontier Science Program (grant no. RGY0056/2022) and the Secretaria d’Universitats i Recerca del Departament d’Economia i Coneixement de la Generalitat de Catalunya (AGAUR 2021-SGR00420). We also thank Centro de Supercomputación de Galicia (CESGA) and the HPC Drago from the Centro Superior de Investigaciones Científicas for access to computer resources.

